## Supplementary_Figures for "CRISPR-finder: A high throughput and cost effective method for identifying successfully edited *A. thaliana* individuals"

```
#ID  Seq  R2
A1F1 CAGTCGAAT ACTCACAGA
A1F2 TCAGTCGAA CTCACAGAT
A1F3 CTCAGTCGA TCACAGATG
A1F4 ACTCAGTCG TGCAGATGA
A1F5 CAGTCGAAT TCACAGATG
A1F6 TCAGTCGAA ACTCACAGA
```

**Supplementary Figure 1:** Example of the text file that is needed for the Plexseq to demultiplex the raw reads based on the frame-shifting nucleotides. The data consist of three columns. The first column indicates the name of the output file after demultiplexing. Columns two and three include the first 9 expected nucleotides for each read on the oligonucleotides that were used for the amplification during library preparation.

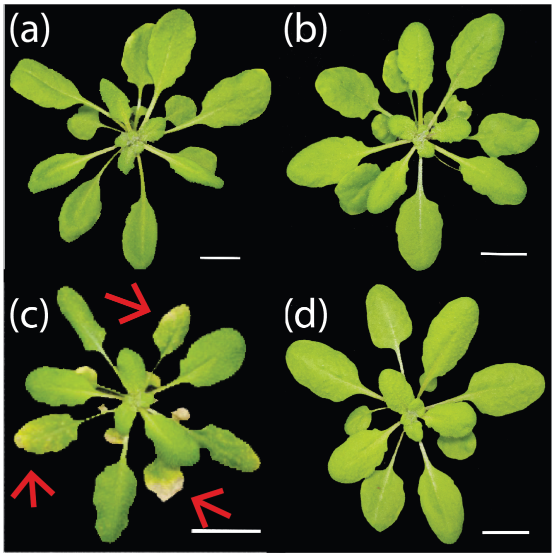


**Supplementary Figure 2:** Morphology of the *ics1/sid2* mutants. The plants were growing in 23°C SD conditions (8 h light/ 16 h dark). The pictures were taken 40 days after sowing. Chlorotic and necrotic regions on leaves are indicated by red arrows. **(a)** Col-0 reference wild type, **(b)** Col-0 *sid2-2* mutant, **(c)** TüWa1-2 wild type, and **(d)** TüWa1-2 *c-ics1-1* mutant. The scale bars indicate 1 cm; note the different sizes of scale bars.
